# Targeting the artemisinin resistant malaria by repositioning of the anti-Hepatitis C Virus drug Alisporivir

**DOI:** 10.1101/2021.01.08.426017

**Authors:** Ayushi Chaurasiya, Swati Garg, Zill e Anam, Geeta Kumari, Nishant Joshi, Jyoti Kumari, Jhalak Singhal, Niharika Singh, Shikha Kaushik, Amandeep Kaur, Neha Dubey, Pallavi Srivastava, Manisha Marothia, Mukesh Kumar, Gobardhan Das, Souvik Bhattacharjee, Shailja Singh, Anand Ranganathan

## Abstract

The rapid emergence of *P. falciparum-resistant* strains raises an urgent need to find new antimalarial drug candidates. This study reports the rational repositioning of the anti-Hepatitis C Virus drug, Alisporivir, a non-immunosuppressive analog of cyclosporin A (CsA) against multiple, drug-resistant strains of *P. falciparum*. Alisporivir being non-hemolytic has been proven to be a better drug than CsA. Indeed, our study also demonstrated the same. Alisporivir inhibited chloroquine-sensitive parasite growth with an IC_50_ of 196.6nM. Alisporivir also inhibited the growth of chloroquine-resistant parasites with an IC_50_ of 422.1nM. Alisporivir exhibited, anti-malarial activity in *in vivo*. Further, we exploited the Cyclophilins targeting potential of Alisporivir against artemisinin-resistant malaria parasite owing to the fact that PfCyP-19B is one of the genes that is overexpressed in artemisinin-resistant parasite revealed by a population transcriptomic study. Our semiquantitative real-time transcript and immunofluorescence analysis confirmed the overexpression of PfCyP-19B in Artemisinin-resistant *P. falciparum* (PfKelch13^R539T^). Artemisinin resistance is attributed to slow clearance of ring stage parasites. Ring survival assay (RSA) is designed to access the potency of compounds on these dormant slow clearing parasites leading to drug resistance. Thus, the potency of Alisporivir against PfKelch13^R539T^ was evaluated by RSA. A 2.5-fold decrease in parasite survival was detected with Alisporivir. Further, combination of Alisporivir with DHA found to potentiate the efficacy of DHA by 4.55-fold. These results support the hypothesis that targeting of resistance mechanism is a potential approach to deal with resistant parasite. Overall, this study demonstrates the rational reposition of Alisporivir against resistant malaria resistance.

## BACKGROUND

Malaria is a vector borne life-threatening disease caused by obligate intracellular *Plasmodium spp*. parasites with an estimated 0.4 million deaths in 2018 (1). *Plasmodium falciparum* has developed drug resistance to all the antimalarial therapeutics leading to demise of multiple first-line treatments, including chloroquine, proguanil, pyrimethamine, sulfadoxine-pyrimethamine, mefloquine, etc. More recently, reports of emerging resistance against Artemisinin-based combination therapies threaten the ongoing global efforts to eliminate malaria and raises alarm for discovering novel therapeutics with potentially new mechanism of action (2).

Artemisinin therapy causes immediate and rapid clearance of ring stage parasites resulting in faster clinical response. However, the short half-life of artemisinin in the blood encourages to discover novel partner drugs that can maintain drug pressure for prolonged duration (3). A number of pharmacologically active compounds have been tested, in combination with artemisinin, on parasite resistant and sensitive strains to access their efficacy in killing parasites. Cyclosporine A (CsA), a known inhibitor of cyclophilins, had been demonstrated to potentiate the action of artemisinin in the early ring stages (4). Thus, it can be used as artemisinin partner drug against artemisinin resistant parasites. However, because of the immunosuppressive nature of CsA, due to its interaction with human calcineurin, and its ability to cause eryptosis, CsA has never been approved as an antimalarial drug (5, 6).

A recently developed CsA analogue, Alisporivir (also known as Debio-025 or DEB025, (D-MeAla3–EtVal4) cyclosporin), remains the most promising second generation cyclophilin inhibitor till date (7, 8). Alisporivir is effective for the treatment of chronic hepatitis C (HCV) infections, HIV, EAV, as well as Duchenne muscular dystrophy (9, 10). It differs slightly from the parent cyclic undecapeptide, in that the sarcosine at position 3 is replaced with Me-alanine, leucine at position 4 is replaced with valine, and the nitrogen at position X is N-ethylated instead of being N-methylated. These chemical modifications not only enhance Alisporivir’s binding affinity for cyclophilins, they also abolish its binding to calcineurin, leading to cessation of immunosuppressive activity (9). Furthermore, Alisporivir has a plasma half-life of 60-90h (10), thus making it a perfect choice for artemisinin partner drug.

This study reports the repositioning of the anti-Hepatitis C Virus drug Alisporivir against multiple, drug-resistant strains of *P. falciparum*. Alisporivir, a non-immunosuppressive analog of cyclosporin A (CsA), exhibits, a potent antiparasitic activity against malaria parasite both *in vitro* culture and *in vivo*, mice model. This drug also targets Artemisinin-resistant strain and potently inhibits the ring-stage survival of Artemisinin-resistant *P. falciparum*. Mechanistically the potent effect of Alisporivir on Artemisinin-resistant *P. falciparum* strain can be explained through its specific targeting of up-regulated PfCyP-19B in resistant strain. Additionally, Alisporivir treatment enhanced the efficacy of DHA against parasite growth suggesting that targeting the resistance mechanism of the parasite is an effective approach to deal with the rapid emergence of drug resistant *P. falciparum*.

## RESULTS

### *In-silico* interaction study of *Pf*CyP-19B with Alisporivir and Cyclosporin A

Cyclophilins (Cyp) are ubiquitous cellular proteins that, in addition to acting as chaperons, possess peptidyl-proline isomerase (PPIase) activity (11). The role of malaria parasite encoded cyclophilins is largely unknown but they represent a potential class of antimalarial drug targets. To evaluate the binding of parasite Cyclophilin with Aliporivir, we performed *in-silico* docking studies of Alisporivir with *Pf*CyP-19B. The structure-refined model of *Pf*CyP-19B having an RMSD score of <2.0 was used for docking with Alisporivir and CsA. The molecular docking results revealed that Alisporivir could efficiently bind to *Pf*CyP-19B with a minimum binding energy of −5.72 (Fig. 1A). Moreover, Alisporivir forms more number of hydrogen bonds with *Pf*CyP-19B compared to CsA (Fig. 1B). Interaction analysis of Alisporivir with amino acid residues of *Pf*CyP-19B demonstrated strong binding of Alisporivir to the protein (Fig. S1).

**FIG 1.**
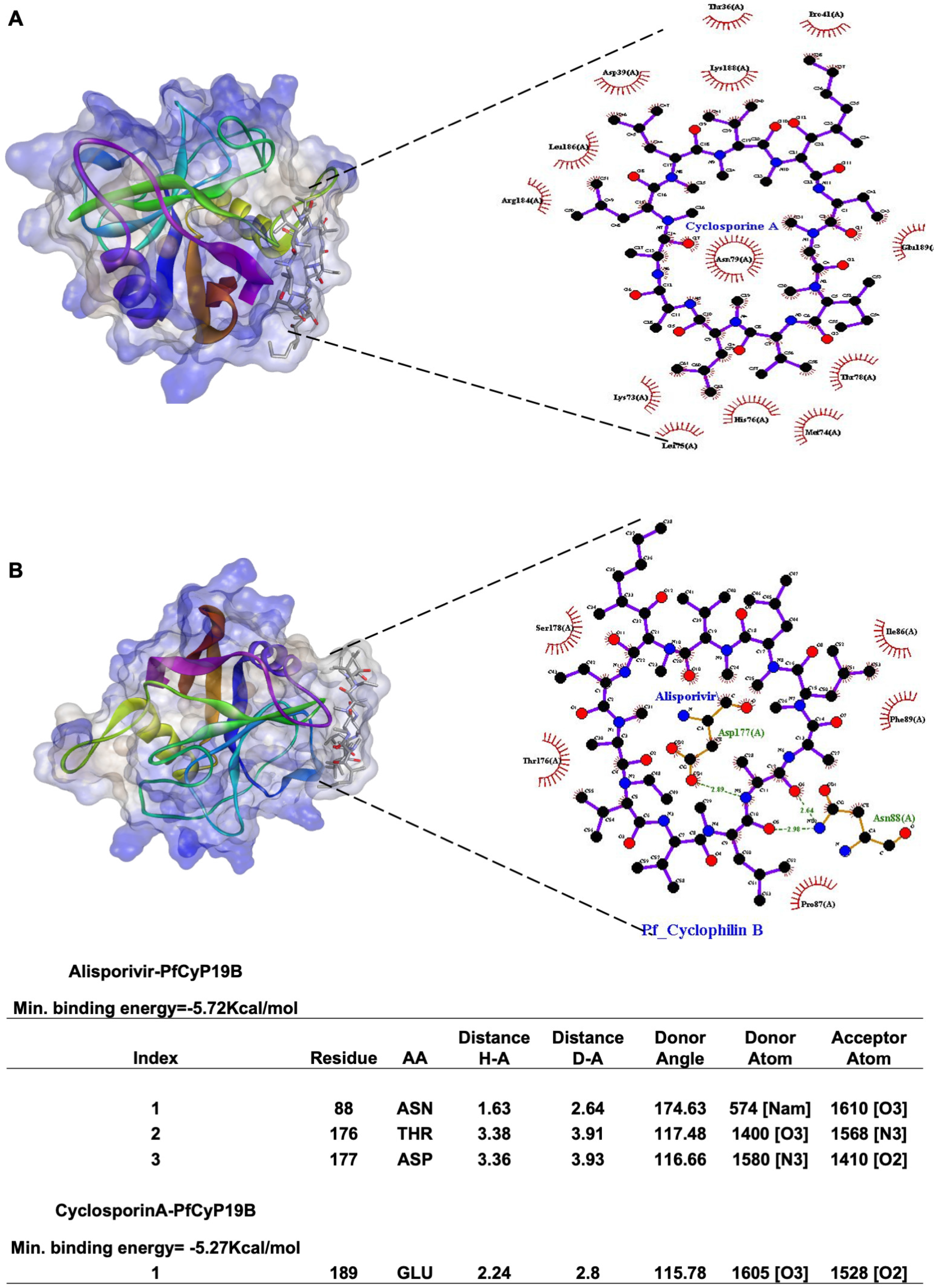
*In-silico* interaction study of PfCyP-19B with Alisporivir and Cyclosporin A. (A). The 3D surface model of CyclosporinA-PfCyP19B complex with ligplot analysis of drug and protein. (B). The 3D surface model of Alisporivir-PfCyP19B complex with ligplot analysis of drug and protein. Details of hydrogen binding residues of CyclosporinA-PfCyP19B and Alisporivir-PfCyP19B complex.

### *Pf*CyP-19B-Alisporivir contacts during dynamic simulation

In order to confirm binding modes and interaction of Alisporivir, the molecular dynamic simulation (MDS) was performed. The behaviour of the *Pf*CyP-19B and Alisporivir was studied in terms of root mean square deviation (RMSD) and root mean square fluctuation (RMSF) between protein, ligand and protein-ligand interactions throughout the simulation for 50 ns. As indicated in Fig. 2A, the Cα and backbone RMSD values of the docked complex showed deviation within the acceptable range of 0-2.7 Å. Also, Lig fit Prot and Lig fit Lig RMSD deviation were in range of 0-2.5 Å. Thus, the presence of the docked molecules did not affect the protein chain conformation. Fig. 2B, indicates areas of the protein that fluctuate the most during the simulation. Typically, the tails (*N*- and *C*-terminal) fluctuate more than any other part of the protein. Secondary structure elements like alpha helices and beta strands are usually more rigid than the unstructured part of the protein, and thus fluctuate less than the loop regions (Fig. S2). Specific *Pf*CyP-19B interactions with the Alisporivir were monitored throughout the simulation. Protein-ligand interactions (or ‘contacts’) were categorized into four types: Hydrogen bonds, Hydrophobic, Ionic and Water bridges and summarized, as shown in the plot (Fig. 2C). Fig. 2D shows timeline representation of the interactions and contacts (H-bonds, Hydrophobic, Ionic, Water bridges) summarized in the Fig 2C. The top panel shows the total number of specific contacts the protein makes with the ligand over the course of the trajectory. The bottom panel shows which residues interact with the ligand in each trajectory frame. Some residues make more than one specific contact with the ligand, which is represented by a darker shade of orange, according to the scale to the right of the plot (Fig. S3 and S4). Apart from that, almost all candidates showed interactions with the crucial amino acids, i.e., GLN70 and ASN109 that occur 64% and 93% of the simulation time, respectively shown in Fig. 2E. It is also observed that crucial candidates have formed more than four interactions and these interactions are responsible for showing the stable behaviour of the complex (Fig. S5).

**FIG 2.**
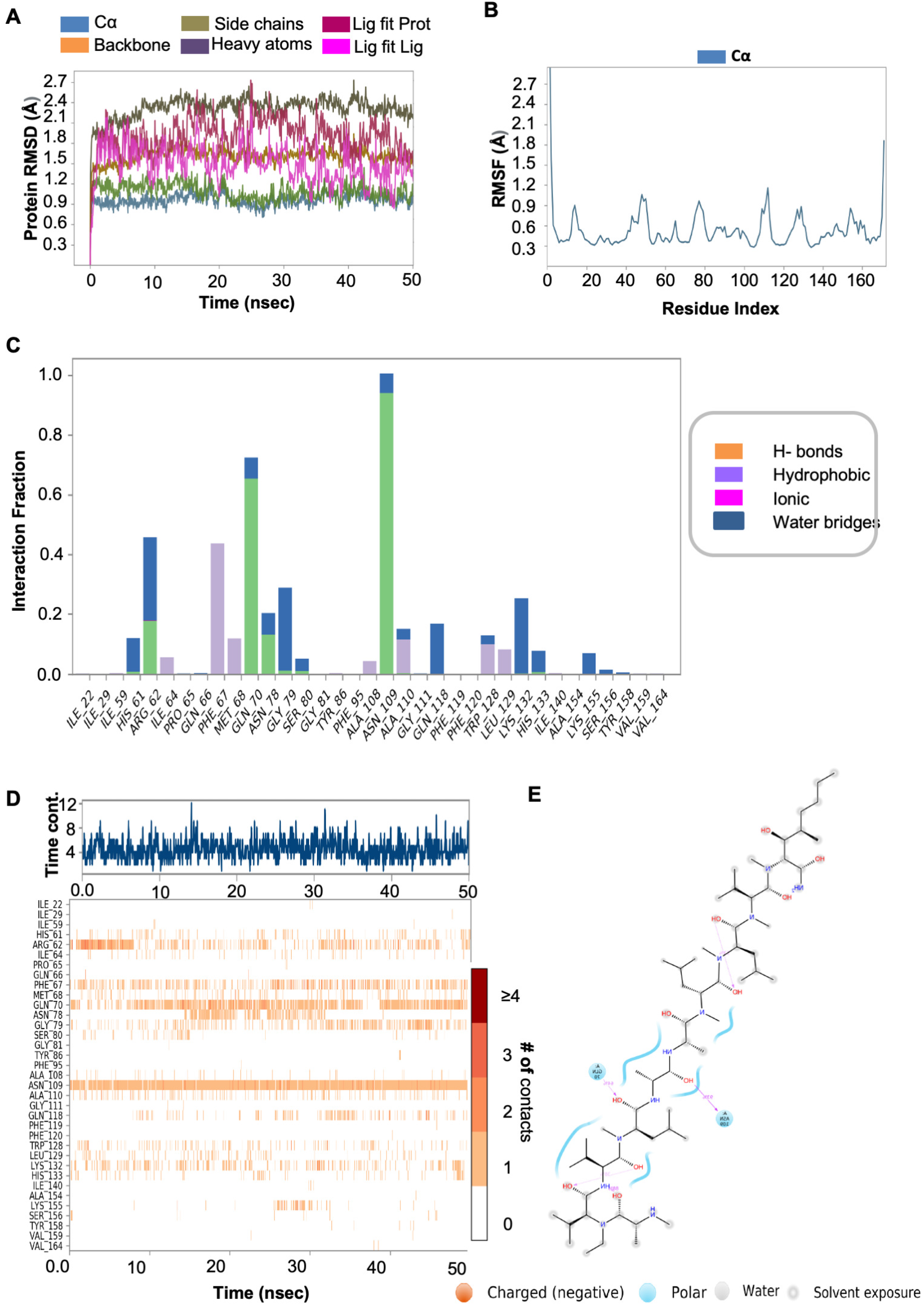
*Pf*CyP-19B-Alisporivir complex RMSD, RMSF and contacts. (A) RMSD of simulated complex. (B) RMSF of simulated complex. (C) Protein-ligand interactions of the MD simulation. All residues that interact with the ligand are shown in the histogram along with the fraction of the simulation for each type of interaction. (D) Analysis of total contacts formed between *Pf*CyP-19B residues and Alisporivir during MD simulation. (E) Analysis of percentage occupancy of the direct interaction between Alisporivir and *Pf*CyP-19B residues.

### Expression analysis of *Pf*CyP-19B in *P. falciparum* 3D7 and R539T strains

Previous reports indicate up-regulation of *Pf*CyP-19B in artemisinin resistant field strains (12, 13). We investigated *Pf*CyP-19B expression level in *P. falciparum* 3D7 and Artemisinin-resistant strain (PfKelch13^R539T^) by RT-PCR analysis (Fig 3A). Increased expression of *Pf*CyP-19B was observed in the Artemisinin resistant strain of malaria parasite which is similar to the earlier published transcriptomics data (12, 13). We next assessed the expression of *Pf*CyP-19B at protein level by immunofluorescence assay (Fig 3B). Expression and localization of *Pf*CyP-19B was investigated in parasite-infected RBCs through confocal microscopy. The staining of *Pf*CyP-19B can be detected in wild type malaria parasites as suggested by green fluorescent spots. The intensity of green fluorescent spots in artemisinin resistant parasites was found to be very high as compared to wild type parasites. Parasite nuclei were stained with DAPI. This indicates the elevated expression of *Pf*CyP-19B in artemisinin resistant parasites as compared to wild type parasites. Therefore, the specific targeting of *Pf*CyP-19B by Alisporivir can combat artemisinin resistance in *P. falciparum*.

**FIG 3.**
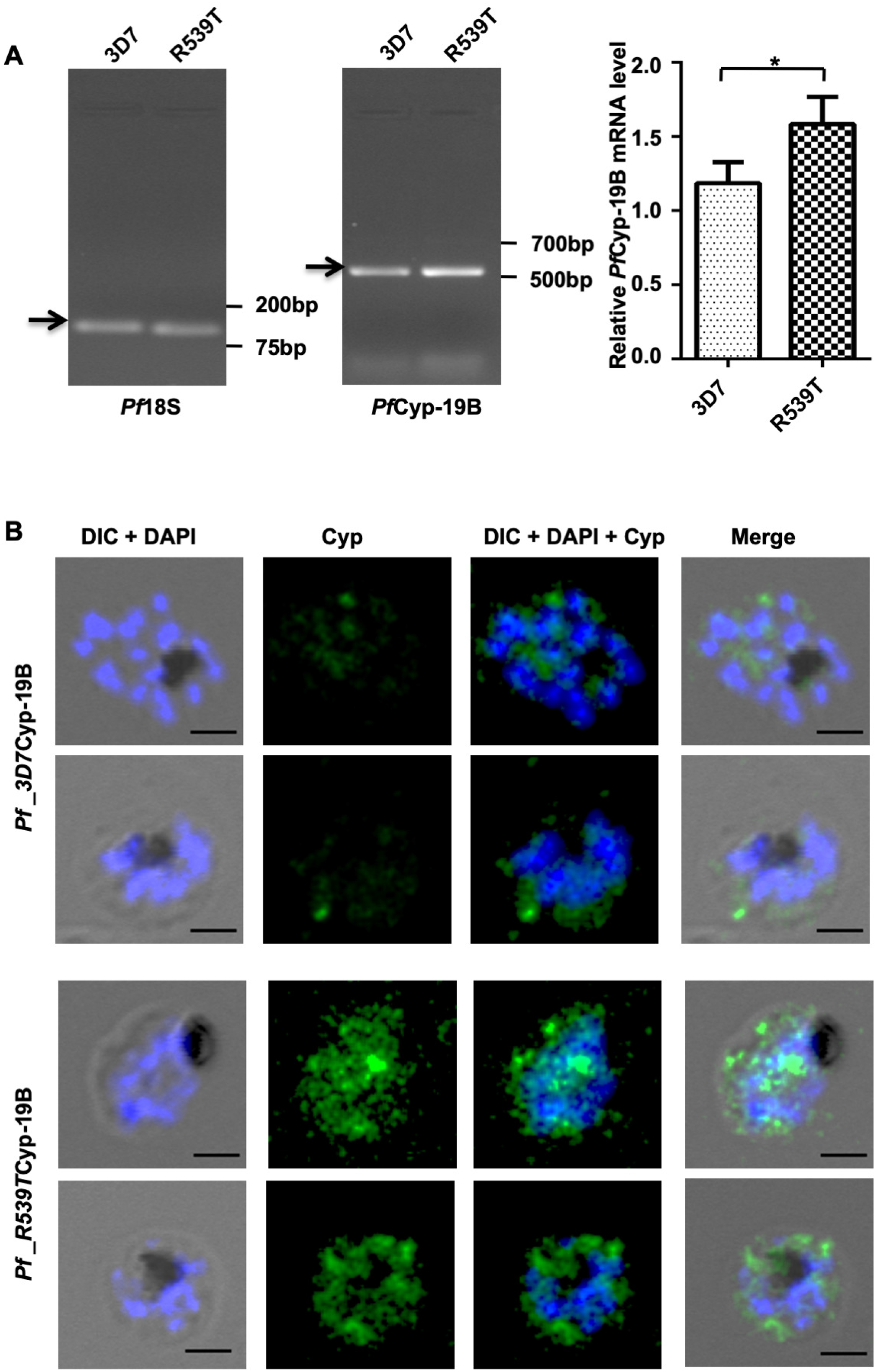
Expression analysis of *Pf*CyP-19B in *P. falciparum* 3D7 and artemisinin resistant R539T strain. (A) Representative image of RNA expression level of *Pf*CyP-19B in *P. falciparum* 3D7 and R539T strains. Quantification of changes in *Pf*CyP-19B RNA expression level was calculated relative to the RNA level of *Pf*18S (internal control). Data represents Mean ± standard deviation of three independent experiments. Statistical significance of difference in RNA level was calculated using the unpaired t-test with Welch’s correction, *P<0.05. (B) Confocal microscopic imaging to evaluate expression pattern of *Pf*CyP-19B protein in parasitized *P. falciparum* 3D7 and R539T strains. Scale bar represents 5 μm.

### *In vitro* anti-malarial activity of Alisporivir against *P. falciparum*

The anti-malarial potential of Alisporivir was investigated by performing growth inhibition assays (GIA) against *P. falciparum* 3D7 strain. Results, assessed by Giemsa staining, indicate that Alisporivir inhibits parasite growth in a dose-dependent manner with an IC_50_ value of 196.6nM compared to 268.5nM for CsA (Fig. 4A). Furthermore, Alisporivir and CsA potently inhibited the growth of chloroquine resistant RKL-9 strain with an IC_50_ of 422.1nM and 439.6nM respectively (Fig. 4B). Morphology assessment of the Giemsa stained parasites was done under the microscope 48 h post-treatment to evaluate the development of parasite. Alisporivir or CsA treated *P. falciparum* 3D7 and RKL strains could not develop into healthy trophozoites and instead demonstrate formation of pycnotic bodies.

**FIG 4.**
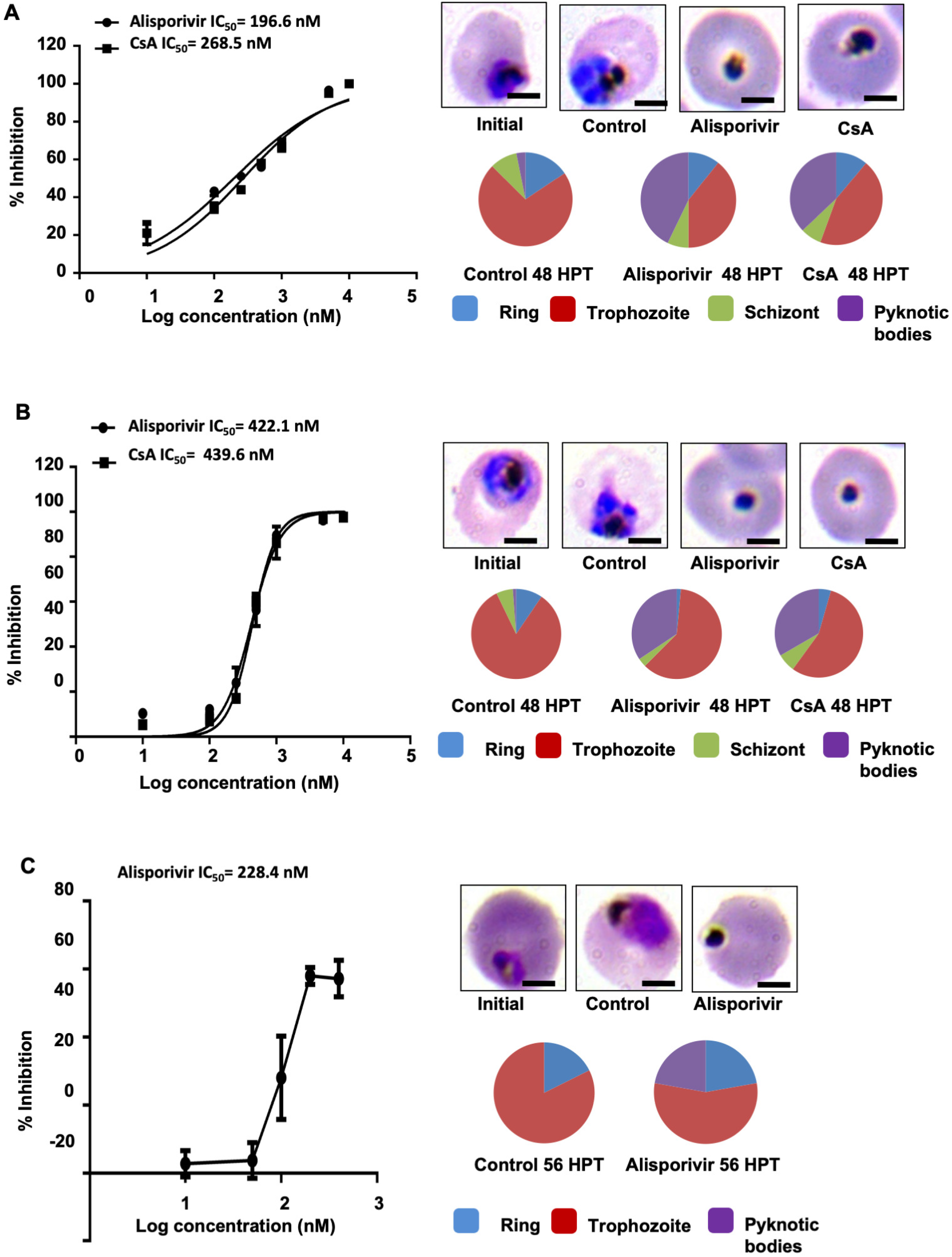
Effect of the Alisporivir on *P. falciparum* growth *in vitro.* Dose-dependent inhibition of parasite growth was monitored in the presence of Alisporivir and CsA (A) *P. falciparum* 3D7 strain (B) *P. falciparum* RKL strain. IC_50_ values of the drugs were determined by plotting the values of percent growth inhibition against log concentrations of drug and calculated by non-linear regression analysis on GraphPad Prism-6 software. Pie charts show relative distribution of different stages of parasites and pycnotic bodies formation at 48 h post infection. (C) Determination of inhibitory potential of Alisporivir against *P. falciparum* artemisinin-resistant strain R539T. IC_50_ value was calculated by GraphPad Prism6 software. Light microscopic examination of parasites morphology after treatment.

The increased expression of *Pf*CyP-19B in artemisinin resistant parasites suggests that it plays an important role in development of drug resistance against Artemisinin. We next examined the antimalarial activity of Alisporivir against Artemisinin-resistant strain (PfKelch13^R539T^) through parasite growth inhibition assay (GIA) at varying concentrations (10 nM-400 nM) of Alisporivir. Alisporivir was found to be potent against artemisinin resistant parasites, with an IC_50_ value of 228.4 nM which is similar to the IC_50_ of wild type parasites. Stage-specific analysis of the morphology of treated parasites at IC_50_ concentration of Alisporivir again demonstrated that treated parasites were reduced to pycnotic bodies while control parasites progressed normally (Fig. 4C).

### Effect of Alisporivir on the eryptosis induction of human erythrocytes

The parent molecule CsA induces death of erythrocytes (5). Therefore, we next investigated the safety of Alisporivir towards erythrocytes. Erythrocytes were incubated with Alisporivir or CsA (10 μM) for 48 h, stained with FITC-annexin-V and assessed by flow cytometry and confocal microscopy. Alisporivir treated erythrocytes did not show any green fluorescence while it was observed on the surface of CsA-treated erythrocytes indicating the exposure of phosphatidyl serine on surface. The number of annexin-V binding erythrocytes was found to be extremely low upon Alisporivir treatment similar to untreated erythrocytes; however, CsA exposure significantly increased the annexin-V positive erythrocytes (Fig. 5A) suggesting that Alisporivir does not induce eryptosis. Next, we investigated possible hemolysis of erythrocytes following Alisporivir treatment.

**FIG 5.**
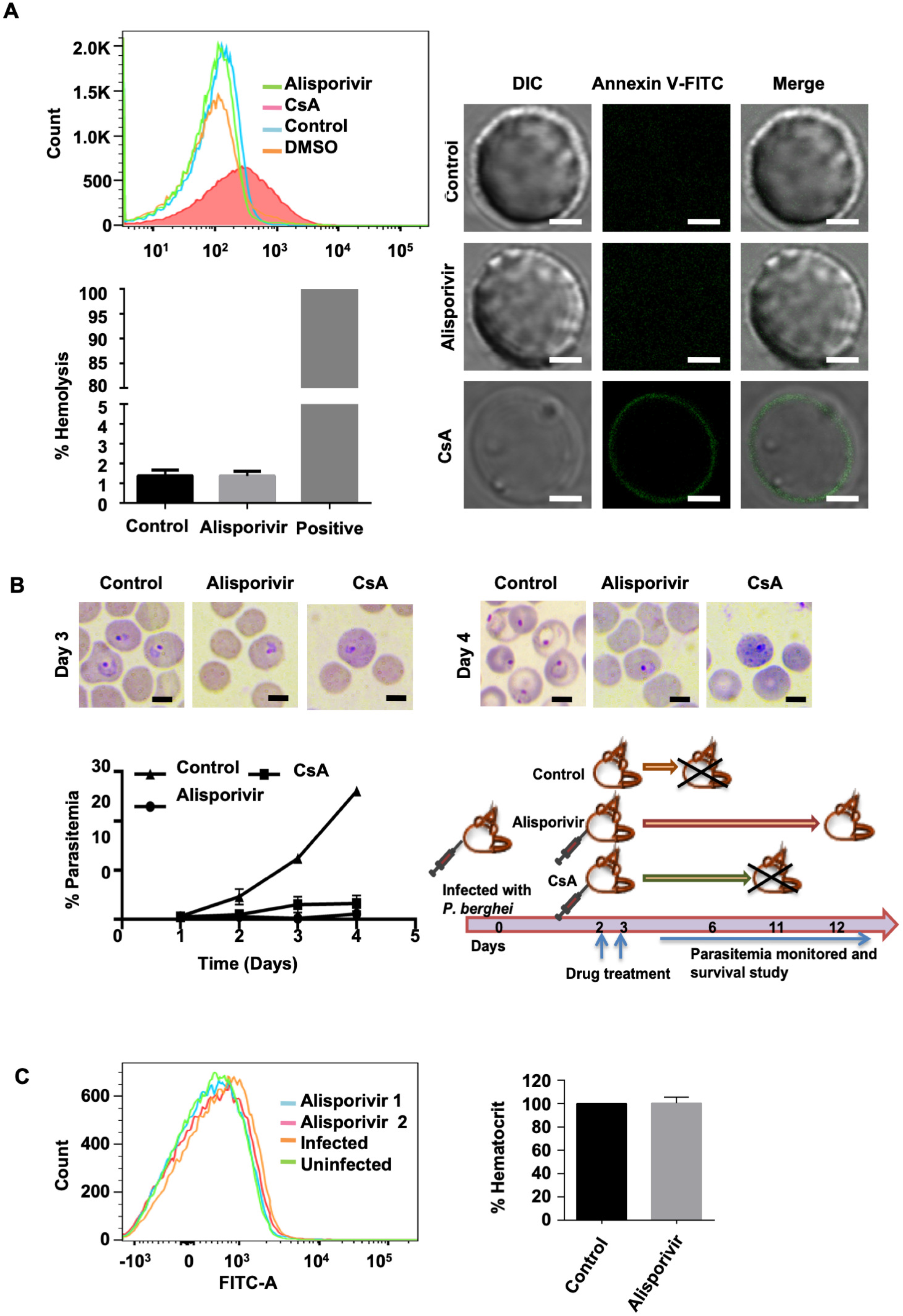
**(A)** Evaluation of eryptosis in human erythrocytes treated with 10 μM of Alisporivir or CsA for 48 h at 37°C using confocal microscopy and Flow cytometry. Representative histogram of annexin-V-binding of erythrocytes following exposure to Alisporivir or CsA. Percent haemolysis was calculated at 10 μM concentration of Alisporivir. Hemolysis value of distilled water diluted RBCs was taken as 100%. (B) Determination of parasitemia in *P. berghei* infected mice with or without treatment of Alisporivir or Cyclosporin A (5mg/Kg b.w). Data represents Mean ± Standard deviation. (C) Representative histogram of annexin-V binding in erythrocytes from *P. berghei*-infected mouse with or without treatment with Alisporivir. Hematocrit percentage in Alisporivir treated mice compared to control. Hematocrit value of control was considered as 100 %. Both groups have similar packed cell volume. Scale bar represents 5 μm.

To measure hemolysis, hemoglobin release from erythrocytes into the media was assessed following 24 h incubation with 10 μM Alisporivir and found it remains unaltered. This observation points towards the fact that Alisporivir is not toxic to the erythrocytes (Fig. 5A).

### *In vivo* anti-malarial efficacy of Alisporivir against *P. berghei*

After confirming the efficacy of Aliporivir in vitro, we studied the antimalarial potency of Alisporivir *in vivo* on mouse malaria parasites, *P. berghei*. Alisporivir and CsA were injected at 5 mg kg^-1^ body weight in *P. berghei* infected mice for two days. The parasitemia in untreated mice, increased with time, up to 25% within 5 days of infection. However, in Alisporivir treated mice we could not find a significant increase in parasitemia. CsA treatment prolonged the prepatent period in line with previous studies (14, 15). Crucially, Alisporivir treatment increased the survival of *P. berghei* infected mice. The untreated animals died within 6 days of infection. In contrast, most of the Alisporivir treated mice survived the infection for more than 12 days (Fig. 5B). In order to study whether Alisporivir caused eryptosis in *P. berghei* infected erythrocytes, the annexin-V-binding property of erythrocytes was evaluated through flow cytometry. Alisporivir treated erythrocytes did not show any significant change in annexin V-binding property compared to uninfected erythrocytes. Percent hematocrit was also found to be similar in infected mice compared to Alisporivir treated infected mice (Fig. 5C).

### Effect of Alisporivir on progression of *P. falciparum* asexual blood stages

To investigate the effect of Alisporivir on progression of asexual parasite blood stages, schizont stage parasite culture of *P. falciparum* 3D7 strain was treated with Alisporivir or CsA (125 nM, 250 nM) and evaluated for 72 h, till the ring stage of third cycle. At each stage, development of parasite was monitored by observing Giemsa-stained smears. Fold change in parasitemia remarkably decreased at the end of the third cycle in treated culture compared to control. As expected, a significant increase in the formation of pycnotic bodies was observed in treated parasite culture (Fig. 6A).

**FIG 6.**
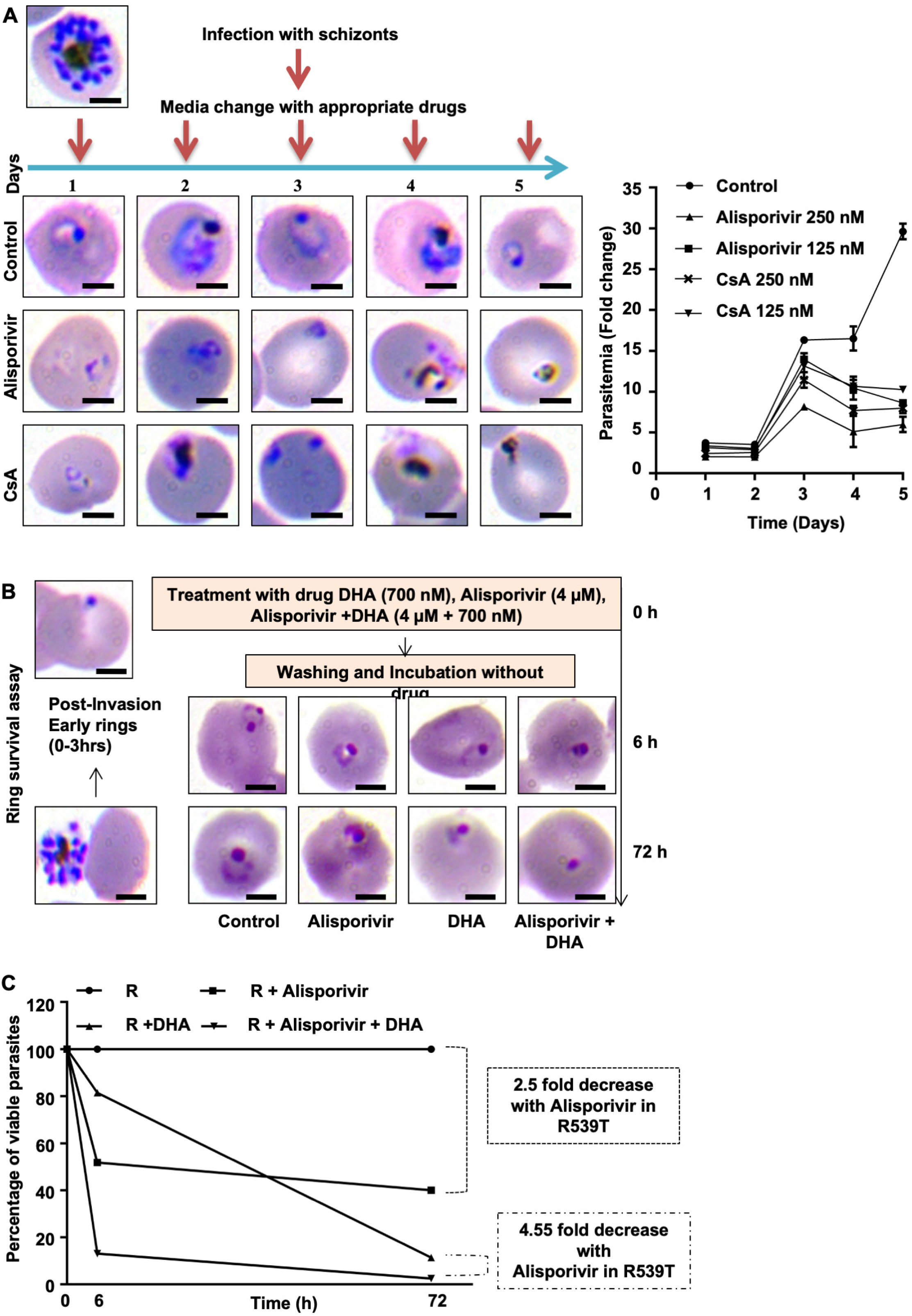
Effect of Alisporivir on progression of *P. falciparum* asexual blood stages and *in vivo* antimalarial efficacy of Alisporivir. (A) Light microscopy images of Alisporivir or Cyclosporine A (250 nM) treated *P. falciparum* parasites. Fold change in parasitemia at different time point of progression. Significant reduction was observed compared to control at day 5 (Alisporivir or CsA 125μM, *p<0.05; Alisporivir or CsA 250 μM, ** p<0.01). Pie diagrams showing relative proportion of different stages of parasites and pycnotic bodies. Alisporivir potentiates antimalarial efficacy of DHA against Artemisinin-resistant *P. falciparum*. (B) Light microscopic evaluation of parasite development following treatment with DHA (700 nM), or Alisporivir (4 μM), or DHA + Alisporivir (700nM+ 4μM) for 6 h. (C) Combination of Alisporivir with DHA enhances antiparasitic activity of DHA against artemisinin-resistant strain (**p<0.01). Tightly synchronized ring stage (0-3 h) culture was used to perform ring survival assay with DHA, Alisporivir or DHA+Alisporivir. Scale bar represents 5 μm. Statistical significance was calculated using the unpaired t-test.

### Alisporivir potentiates antimalarial efficacy of DHA against Artemisinin-resistant *Plasmodium falciparum*

Artemisinin resistance is attributed to slow clearance of ring stage parasites. Ring survival assay (RSA) was designed to access the presence of these dormant slow clearing parasites leading to drug resistance. To determine the potency of Alisporivir against Artemisinin-resistant parasites, ring survival assay (RSA) was performed on PfKelch13^R539T^ strain with DHA, Alisporivir, or a combination of DHA with Alisporivir. Tightly synchronized ring stage culture was treated with the drugs for 6 h. Following treatment, culture was maintained without drug pressure to determine parasite viability. Ring stage development of parasites was observed to study the effect of drug exposure after 6 h and 72 h of post-treatment. Control parasites could progress to form healthy ring stage parasites while in DHA and Alisporivir treated cultures, dead parasites could be observed (Fig. 6B). After both 6 h and 72 h of post-treatment, Alisporivir and DHA exhibit decreased parasite survival in the artemisinin resistant strain. Parasite survival was decreased by 2.5 fold after treatment with Alisporivir. Interestingly, combination of Alisporivir with DHA further potentiated the efficacy of DHA by 4.55 fold in resistant strain after 72 h of treatment and decreased the parasite survival at the early ring stage (Fig. 6C).

Taken together, *Pf*CyP-19B appears to be involved in Artemisinin-resistance; and our results exploit the possible mechanism that contributes to drug resistance by targeting *Pf*CyP-19B with Alisporivir.

## DISCUSSION

The unavailability of new potent antimalarials and emerging resistance against Artemisinin and its partner drugs are the major obstacles in malaria control (2). Therefore, there is an urgent need for identification of novel drug targets and antimalarial agents to prevent occurrence of drug resistance. Repositioning of existing drugs as novel antimalarials offers a new way for drug development (16, 17). In this study, we have repositioned an anti-Hepatitis C Virus drug Alisporivir and demonstrate its potency against multiple drug-resistant strains of *P. falciparum* and mouse malaria in vivo. Importantly, Alisporivir does not cause eryptosis which is a common side effect of the parent molecule CsA We further demonstrate that Alisporivir targets overexpressed *Pf*CyP-19B in artemisinin resistant strain and strongly binds with *Pf*CyP-19B in *in-silico* studies. More importantly, in ring survival assay, combination of Alisporivir with DHA potently inhibits survival of the Artemisinin-resistant parasites indicating important role of *Pf*CyP-19B in the mechanism of Artemisinin-resistance development. Altogether, our data demonstrate that Alisporivir can be used as an effective antimalarial or as a partner drug with Artemisinin for malaria treatment.

Fast-acting antimalarial drugs have revolutionized the malaria treatment paradigm with huge success rates for all parasite strains. However, challenges still remain in certain populations that have developed resistance and experienced treatment failure. To cure *Plasmodium spp*. infection, a treatment regimen must combine fast acting antimalarial potency of a drug along with a combination drug having higher plasma life. This will ensure killing of residual parasites and detain the development of drug resistance. Alisporivir, with an estimated plasma half-life in range of 60-90h fulfills this need and hence is suitable for once-daily administration (9, 10). In addition, clinical pharmacokinetic studies have shown rapid absorption of Alisporivir after oral administration (<2 hours) and high plasma accumulation of Alisporivir after four weeks of medication. Alisporivir has shown an acceptable clinical safety profile in clinical studies of HCV patients to date (9, 10).

Cyclophilins, the target of Alisporivir, are present in all domains of life and are crucial for protein folding (18). In *Plasmodium spp*. cyclophilins represent an important drug target (19). Our previous report also has identified and confirmed the human CyP-B as a novel druggable target (20). In our study, we found that the compound Alisporivir is effectively docking around the binding site in PfCyP-19B with minimal free binding energy. Furthermore, in our study, 50 ns molecular dynamic simulation revealed that the observed deviation values of docked complexes were found to be less or equal in the range of the reference protein, i.e. the first simulation frame. This specifies that the complexes were stable during the simulation of molecular dynamics, and no significant conformational changes were observed. Overall, the *in-silico* data indicates, that the Alisporivir stably docks well within the binding site of the *Pf*CyP-19B protein.

Furthermore, the observation that expression of *Pf*CyP-19B upregulates in the artemisinin resistant field parasites opens new avenues for targeting it and combat drug resistance malaria (12, 13). Our data confirms the increased expression of *Pf*CyP-19B transcript as well as protein in artemisinin resistant *P. falciparum* parasites. And, the *in-silico* results demonstrate the strong binding of Alisporivir with *Pf*CyP-19B encouraged us to explore Alisporivir in detail for its prospective antimalarial properties. Alisporivir demonstrates potent antimalarial activity against chloroquine sensitive (196.6 nM), chloroquine resistant (422.1 nM) and artemisinin resistant (228.4 nM) strains of *P. falciparum*. In addition, Alisporivir shows efficacy against *P. berghei in vivo* mouse model. Notably, the two-dose Alisporivir treatment reduces *P. berghei* load in mice and increase the survival of host. Understanding the mechanistic importance of *Pf*CyP-19B overexpression in artemisinin resistance and its specific targeting by Alisporivir offers a novel approach for drug development.

The secret of artemisinin resistant parasites is their potential to remain dormant under drug pressure but as the drug levels in plasma goes down, notably the elimination half life of artemisinin in plasma is 2-5h, the parasites recover from the artemisinin exposed dormancy (21). Therefore the conventional 48h assay cannot uncover the presence of drug resistant parasites. Hence we performed RSA to identify the presence of resistant parasites in the culture and demonstrate that Alisporivir in combination with DHA potently reduces ring-stage survival of the Artemisinin-resistant parasites. This data highlights the efficacy of Alisporivir against drug resistant parasites and fulfills the requirement of novel drugs for drug resistant malaria.

Earlier studies have demonstrated that CsA is an effective antimalarial against multiple strains of parasite, but due to its immunosuppressive and eryptosis inducing nature could not be pursued further for malaria drug development (4, 5, 14, 15, 20, 22, 23). However, we demonstrate that Alisporivir, which is a non-immunosuppressive drug, does not induce eryptosis both *in vitro* (human erythrocytes) and *in vivo* (mouse erythrocytes). Our data and previous studies thus validate Alisporivir as a safe drug which can be taken forward for antimalarial drug development.

In a previous study, Tong *et al* demonstrated that cyclophilin targeting drug CsA in combination with Artesunate have a synergistic anti-malarial activity against artemisinin resistant strain IPC 5202 in ring-stage survival assay. The combination of CsA with Artesunate displayed sum FIC_50_ value of 0.679 ± 0.017, thus suggesting CsA could be a potential drug partner with AS (4). These results further support our findings for Alisporivir, which is a non-immunosuppressive analog of CsA.

In conclusion, the identification of the potent inhibitory effect of Alisporivir on malaria parasite growth, its functioning with minimal side effects, open the immediate possibility of its use, either alone or in combination with existing antimalarials, as a potent candidate drug against malaria parasites.

## METHODS

### Chemicals

Alisporivir (DEB-025; Debio-025, MedChem Express, USA) and CsA (Sigma-Aldrich, USA) stocks were prepared in DMSO.

### Parasite Culture

#### *In vitro* culture of *Plasmodium falciparum*

*P. falciparum* 3D7, RKL9 and PfKelch13^R539T^ strains were maintained at 2% hematocrit in RPMI 1640 medium (Invitrogen, USA) supplemented with 27.2 mgL^-1^ hypoxanthine and 0.5% Albumax I, at 37°C mix gas environment using O^+^ red blood cells (obtained from Rotary blood bank, Delhi), as described previously (3). Culture was synchronized with 5% sorbitol and routinely monitored by Giemsa staining (Sigma, USA). The *P. falciparum* 3D7 transgenic parasite isolates bearing artemisinin resistant mutation PfKelch13^R539T^ (R539T) were used in this study and the mutant parasite line was generated as described previously (24).

#### Infection (*In vivo* proliferation of *Plasmodium berghei*)

Animal studies were approved by the Institutional Animal Ethics Committee (IEAC) of JNU, New Delhi and performed according to CPCSEA guidelines. Mice were obtained from Central Laboratory Animal Resources, JNU, New Delhi and maintained under standard conditions.

*P. berghei* ANKA strain (ATCC, India) parasitized mouse erythrocytes (1×10^5^) were injected intraperitoneally in BALB/c mice. Three groups of mice were injected with CsA (5mg/Kg, intraperitoneal), Alisporivir (5mg/kg, intravenous), and buffer control respectively for two days. Parasitemia was determined from Giemsa stained tail blood smears.

### *In vitro* growth inhibition assays

Synchronized *P. falciparum* parasites at 0.8-1.5% initial parasitemia and 2% hematocrit were treated with Alisporivir or CsA at concentrations ranging from 10 nM-10 μM (3D7 and RKL-9 strain) and 10 nM-400 nM (PfKelch13^R539T^strain); DMSO treated culture was considered as control. Following incubation, parasitemia was determined through counting of Giemsa-stained thin blood smears. More than 2000 RBCs were counted per slide and IC_50_ calculated using graphpad prism 6.0 (CA, USA).

Percent growth inhibition was determined as follows: % Inhibition = [1 - % Parasitemia of treatment/% Parasitemia of Control] * 100. To analyse the progression of drug treated parasite in comparison to control, drug-treated culture was monitored for five days till the ring stage of third cycle. At each stage, parasitemia and morphology of parasites were examined by Giemsa-stained blood smears.

### Flow cytometric analysis of annexin-V binding

To evaluate eryptosis in erythrocytes, flow cytometric analysis was performed, by measuring annexin-V fluorescence intensity. After treatment, the erythrocytes were washed in Annexin-V binding Buffer (140 mM NaCl, 10 mM HEPES (pH 7.4), and 2.5 mM CaCl_2_). Erythrocytes were incubated with annexin-V-FITC (Thermo Scientific, USA) in annexin-V binding buffer (1:20) for 20 min at room temperature. Samples were then analyzed by flow cytometry (BD, LSR Fortessa, USA) using excitation at 488 nm and an emission at 530 nm. Identical procedure was followed in order to measure eryptosis in mice samples.

### Hemolysis assay and hematocrit determination

For the hemolysis assay, RBCs (1x 10^7^) were treated with Alisporivir (10 μM) at 37 °C, 5 % CO_2_ incubator for 24 hours. After incubation, absorbance of supernatant was measured at 415 nm with a microplate reader (Thermo Scientific, USA) to assess the hemoglobin leakage. Erythrocytes lysed in distilled water were used as a positive control. To measure hematocrit, blood was collected from infected and Alisporivir-treated infected mice. Hematocrit was determined as the percentage of blood cells in the total volume of blood.

### Reverse transcription polymerase chain reaction (RT-PCR)

Total RNA of *P. falciparum* 3D7and PfKelch13^R539T^ strain was isolated using TRIZOL (Life Technologies, USA). Equal amount of total RNA was used for cDNA synthesis using the cDNA synthesis Kit (Thermo fisher, USA). RT-PCR was performed using specific primers (Table S1) for 20 cycles of amplification and the expression level of *Pf*CyP-19B gene (PlasmoDB Id: PF3D7_1115600) quantified using Image J software (NIH). *Pf*18S was used as an internal control.

### Immunofluorescence assay

Expression level of *Pf*CyP-19B in *P. falciparum* 3D7 and PfKelch13^R539T^ strains was monitored through the immunofluorescence assay. Schizont-stage *P. falciparum* infected erythrocytes were used to prepare thin smear and subsequently fixed with chilled methanol. Slides were blocked with 3% BSA and stained next with CyP-B antibody (1:200 dilution) for 1 h at RT. Subsequent to washing, the slides were incubated with Alexa 488 conjugated anti-mouse secondary antibody (1:500 dilution, Invitrogen) at RT for 1 h. Mounted slides with DAPI-antifade (Invitrogen, USA) were examined with confocal microscope (Nikon, USA).

### Ring Survival Assay

To assess the antimalarial potential of Alisporivir with DHA (Sigma-Aldrich, USA), ring survival assay was performed on Artemisinin-resistant strain as described previously (3, 25). Briefly, percoll-purified, early ring-stage parasite culture (0-3 h post-invasion) was treated with Alisporivir (4 μM), DHA (700 nM) and Alisporivir in combination with DHA (4 μM+700 nM) for 6 h followed by washing with complete media and incubated in fresh complete medium without the drug for 66 h. After the drug treatment, at 6 h and after incubation at 72 h, thin blood smears of the samples treated as well as the control were prepared and >10,000 RBCs analysed using Giemsa-staining. Percent parasite survival was determined by comparing the number of viable parasites in treated samples compared to untreated control.

### *In-silico* docking studies

Protein structure files were downloaded from Protein databank. 3D protein structure of plasmodium cyclophilin (PDB ID: 1QNG) was obtained from Protein databank. Chemical structures of compounds were obtained through the Drugbank. Protein and Ligands structures were optimized using Swiss PDB viewer version 4.1.0 and ChemBio3D ultra 12.0 respectively. Autodock version 1.5.7rc1 and Cygwin terminal were utilized to execute the docking commands. Binding site for the ligand was chosen around its catalytic domain residues. Chimera, Ligplus, Discovery Studio version19.1.0 and Pymol version 2.3.2 software were used for further analysis and visualization of docking results (26).

### Molecular dynamics simulation

Molecular dynamics simulation (MDS) was performed on Desmond version 2018.4, running on Linux operating system (27). First, the protein-ligand complexes (crystal structure of *Pf*CyP-19B bound with Alisporivir obtained by docking) were prepared in protein preparation wizard of Desmond module with the explicit solvent model and SPC and orthorhombic box shape. The box size was set at distance of 10 Å from the outermost atom of the protein-ligand complex. Sodium chloride with approximate physiological concentration of 0.15 M was placed to the simulation box to set the ionic strength. After the minimization jobs were performed to relax the system into a local energy minimization, this model system was submitted to MD simulation steps using Desmond package with OPLS_2005 force field. In Maestro, MD simulation was analyzed by the Simulation Event Analysis tool, and ligand-receptor interactions were identified using the Simulation Interaction Diagram tool.

### Statistical analysis

Statistical significance of data was measured using two-tailed Student’s t-test where error bar represent mean ± SD of three independent experiments.

## Acknowledgement

We thank Central Instrumentation Facility (CIF) of Special Centre for Molecular Medicine, JNU. We also thank Advanced Instrumentation and Research Facility (AIRF), JNU, New Delhi for confocal microscopy. This work is supported by Department of Science and Technology, SERB grant number CRG/2019/00223, and IRHPA IPA/2020/000007 (AR, SS). Funding from Science and Engineering Research Board (SERB) (EMR/2016/005644) and Drug and Pharmaceuticals Research Programe (DPRP) (Project No. P/569/2016-1/TDT) for SS is acknowledged. AC is supported by senior research fellowship from department of biotechnology (DBT), New Delhi, India.

## Author Contribution Information

AR and SS conceptualized the idea, supervised and designed the experiments. AC and SG performed the experiments, validated, analysed data and wrote the manuscript. SG, GK, JK set up malarial cultures, conducted experiments and helped in imaging analysis. NJ did bioinformatics analysis. ZA wrote parts of the manuscript. ZA, JS, NS, SK, AK, ND helped in analysis and experimental design. MM, PS, MK helped in experimental methodology. AR, SS, GD, AC, SG critically analyzed the data and the manuscript. SB has generated the Artemisinin resistant strain.

## Additional Information

### Compliance with ethical standards

All the experiments were approved by institutional IBSC committee and performed in accordance with the guidelines and regulations of Jawaharlal Nehru University. Animal handling and experiments were approved by the Institutional Animal Ethics Committee (IAEC), Jawaharlal Nehru University, New Delhi and performed as per CPCSEA guidelines.

## Abbreviations

CsA: Cyclosporin A
CypB: Cyclophilin B
DHA: Dihydroartemisinin

